# The GTPase ARFA1 interactor Cullin 3 Substrate-adaptor Protein 1 (CSP1) positively modulates nodulation

**DOI:** 10.64898/2025.11.30.691430

**Authors:** Carolina Rípodas, Marina Cretton, Andrés Eylenstein, Claudio Rivero, María Eugenia Zanetti, Flavio Blanco

## Abstract

Legume plants have the capacity to incorporate atmospheric nitrogen by establishing an endosymbiotic interaction with soil bacteria resulting in the formation of nitrogen-fixing nodules. Bacteria are internalized through a tightly regulated process that requires membrane remodelling and vesicle trafficking, which are controlled by small GTPases. Members of the ARF family of GTPases mediate vesicle budding in a wide range of biological processes; however, the modulation of ARF members, their subcellular localization and the formation of complexes with other proteins during the root nodule symbiosis has not been investigated. Here, to identify proteins that physically interact with MtARFA1, a yeast two hybrid screening was performed using a cDNA library of *Medicago truncatula* roots inoculated with *Sinorhizobium meliloti*. One of the identified MtARF1 interactors is a protein that possesses a BTB/POZ domain. BTB/POZ domains are present in substrate-specific adaptors that form complexes with the Ubiquitin ligase E3 Cullin3 (CUL3), thus the interactor was designated as *M. truncatula* CUL3 substrate-adaptor protein 1 (MtCSP1). Physical interaction between MtARF1 and MtCSP1 was verified in planta by co-immunopurification assays and bimolecular fluorescence complementation, revealing that the interaction takes place in vesicles of the late endosome. The *MtCSP1* promoter is active in lateral roots and in the meristem of indeterminate nodules. Phenotypic analysis of transgenic roots with altered mRNA levels of *MtCSP1* evidenced the requirement of this gene for the progression of rhizobial infection and nodule organogenesis. This work establishes a link between small GTPases and protein degradation by the ubiquitin system in the context of the nitrogen-fixing symbiosis.

**Significant statement:** Small GTPases are molecular switches required for rhizobial infection in the root-nodule symbiosis; however, little is known about how their levels are regulated during this process. We identified a substrate-adaptor protein that interacts with ARFA1, connecting this monomeric GTPase with protein degradation via ubiquitination during the activation of the genetic programs of symbiosis: rhizobial infection and nodule organogenesis.

## INTRODUCTION

Gene expression is controlled at different regulatory levels, from transcriptional and post-transcriptional levels to the cytoplasmic translational and posttranslational events in the case of protein-coding genes. Protein degradation can dynamically modify the level of a subset of proteins in response to environmental stimuli or as part of developmental programs. The ubiquitin-proteosome system, a protein degradation complex highly conserved in eukaryotes, comprises the action of enzymatic activities that catalyze the ubiquitin activation (E1), conjugation (E2) and ligation (E3). As a result, proteins are conjugated to a chain of ubiquitin residues that target them for degradation. Ligation of ubiquitin is produced by the action of four types of E3 ligases: the monomeric HECT, RING, and U-box, and the multimeric complex Cullin-RING ligases (CRLs). CRLs include a cullin, the adapter protein RBX1 that binds to an E2 ubiquitin conjugating enzyme and substrate adapter proteins (Chen & Hellmann, 2013; Ban & Estelle, 2021). Cullin E3 ligases, which are crucial for many biological processes, interact with Bric-a-Brac Tramtrack and Broad complex/POx virus and Zinc finger (BTB/POZ) substrate adapters (Ban & Estelle, 2021). All these substrate adapters contain one or two BTB/POZ domains that interact with CUL3, and additional protein-protein interaction domains (e.g. armadillo, ankyrin, MATH) that define the substrate specificity (Ban & Estelle, 2021). BTB/POZ domains containing proteins have been shown to participate in development, hormone responses and plant immunity (Chen & Hellmann, 2013; Ban & Estelle, 2021). Noot1 and Noot 2 are two well characterized BTB/POZ proteins that maintain nodule identity by specifying the cell fate of nodule cells (Ferguson & Reid, 2005; Shen *et al*., 2019; Liu *et al*., 2023). However, the biological role of many of these BTB/POZ proteins and their contribution to the specificity of cullin-mediated protein degradation are not well characterized.

Biological nitrogen fixation (BNF) has emerged as a sustainable alternative to alleviate the negative impact caused by the excessive use of chemical fertilizers and optimize crop yields. BNF consists of the reduction of atmospheric N_2_ to ammonium by living organisms, allowing the incorporation of nitrogen into ecosystems. In particular, leguminous plants (*Fabaceae*) establish a symbiotic association with nitrogen-fixing soil bacteria, known as rhizobia. This association allows plants to assimilate nitrogen metabolites, while rhizobia obtain carbon and energy in the form of sugars. The process depends on the formation of specialized root organs known as nodules, where rhizobia are allocated to fix nitrogen. The formation of functional nodules depends on two developmental programs that run in coordination, bacterial infection through cells of the epidermis to reach cortical cells (Murray, 2011), and the nodule organogenesis, which is initiated by reactivation of cell divisions in the cortex, endodermis and pericycle (Xiao *et al*., 2014). In most agronomically important legume crops, infection occurs intracellularly through the root hair. Attachment of rhizobia triggers a change in the direction of the root hair growth, resulting in a curl that entraps rhizobia and forms an infection pocket. A tubular structure, known as the infection thread (IT), is then formed by deposition of membrane and cell material transported by an active vesicle traffic (Murray, 2011). The IT transverses the epidermal cell leading rhizobia toward the cortical cells, where reactivation of cell division occurs to give origin to the nodule primordium. When the IT reaches the primordium, bacteria are internalized into the cells, differentiate into bacteroids and become surrounded by a membrane derived from the plant plasma membrane to form organelle-like subcellular structures called symbiosomes, where the bacteroids carry out the nitrogen fixation reaction (Zhang, X *et al*., 2024).

The formation of ITs requires degradation of the cell wall, polar growth of the invagination and deposition of new material through active intracellular trafficking of vesicles (Murray, 2011). Monomeric GTPases regulate all four stages of this trafficking: formation, transport, recognition and fusion of the vesicle with the target membrane (Rutherford & Moore, 2002; Rivero *et al*., 2017). The plant monomeric GTPases superfamily is subdivided into four main families based on structural and functional similarities: RAB, ARF, RAN and RHO (Rivero *et al*., 2017). Different members of these families of GTPases have been implicated in several stages of symbiosis: Rab7 from *Medicago truncatula* was shown to be required for maturation of the symbiosome (Limpens *et al*., 2009) and two Rab proteins from soybean, Rab1p and Rab7p, are required for the formation of the symbiosome membrane (Cheon *et al*., 1993). Other GTPases belonging to the Rho of plants (ROP) subfamily play roles during early signaling and infection during nodulation in *Lotus japonicum* and *M. truncatula* (Ke *et al*., 2012; Kiirika *et al*., 2012; Lei *et al*., 2015; Ke *et al*., 2016).

In the grain legume *Phaseolus vulgaris,* we have previously described that the gene encoding a RAB of the A2 family (*PvRabA2*) is activated in response to rhizobial strains with high nodulation efficiency (Peltzer Meschini *et al*., 2008). Reverse genetic studies using RNA interference (RNAi) revealed that *PvRabA2* is strictly required for root hair deformation and IT initiation, indicating that morphological changes that precede bacterial infection are dependent on PvRabA2 (Blanco *et al*., 2009). Phenotypic analysis of plants expressing miss-regulated mutant variants of RabA2 (constitutively GTP-bound and constitutively GDP-bound variants) exhibited premature release of rhizobia into the cytoplasm of epidermal cells, suggesting that fine-tune modulation of PvRabA2 activation is required to maintain the integrity of the IT membrane (Dalla Via *et al*., 2017). Subcellular localization assays showed that PvRabA2 is mainly associated with moving vesicles located near the plasma membrane in roots (Blanco *et al*., 2009), specifically in Golgi stacks and the trans-Golgi network (Dalla Via *et al*., 2017). This result was supported by the colocalization of PvRABA2 with ARFA1 (ADP Ribosylation Factor 1), a monomeric GTPase of the ARF family, whose localization in Golgi and trans-Golgi vesicles has been well documented in different organisms (Takeuchi *et al*., 2002; Xu & Scheres, 2005; Stefano *et al*., 2006; Robinson *et al*., 2011).

In this work, we identified a *M. truncatula* protein that physically interacts with MtARFA1 in a yeast two hybrid assay and *in planta*. This MtARFA1 interacting-protein displays high identity with a formin-interacting protein from Arabidopsis that contains a BTB/POZ domain and is predicted to act as a substrate adaptor for protein degradation via the ubiquitin-proteosome pathway in the context of nitrogen-fixing symbiosis.

## RESULTS

### Identification of a BTB/POZ containing-protein that physically interacts with ARFA1

First, we attempted to identify proteins that interact with the monomeric GTPase PvRABA2. A yeast two hybrid screening was performed using PvRABA2 as a bait and a cDNA library from common bean, previously obtained in our lab (Battaglia *et al*., 2014). As monomeric GTPases interact with effector proteins when they are in the GTP-bound conformation, we introduced a point mutation in the RABA2 sequence to obtain a constitutively active (CA) version of the bait. This screening did not yield positive colonies using either the wild type or the CA versions of RABA2. Considering this result, we decided to screen for interactors of ARFA1, another monomeric GTPase that colocalizes with RABA2 (Dalla Via *et al*., 2017) and is strongly coregulated during early stages of symbiosis (Supplemental Figure 1). ARFA1 is a branch of the ARF family composed of several genes that encode nearly identical proteins in different legume and non-legume plants (Rivero *et al*., 2017). Proteins of this subfamily have been shown to be required for root growth in *Arabidopsis thaliana* (Xu & Scheres, 2005). In order to identify proteins that physically interact with MtARFA1, a yeast two hybrid assay was performed using a *M. truncatula* cDNA library fused to the Gal4 activation domain, prepared from total RNA extracted from roots 4 days after inoculation with *S. meliloti* strain 2011, and cloned into the pGAD-HA vector. As bait, we used the open reading frame of the MtARFA1 (MtrunA17_Chr5g0413451) protein with a point mutation at the amino acid 71, where a glutamine was exchanged for a leucine (Q71L). This change affects the GTP hydrolysis activity, resulting in a CA MtARFA1 version. After the screening, we obtained fourteen yeast positive colonies corresponding to two different genes: MtrunA17_Chr4g0019731 that encodes for a small signalling peptide of the DEVIL/ROTUNDIFOLIA (DVL/ROT) family, and MtrunA17_Chr6g0461001, which encodes for a protein with high homology to a formin-interacting protein from Arabidopsis (Banno & Chua, 2000). Analysis of the predicted protein in *M. truncatula* showed the presence of a BTB/POZ domain, which has been reported to form a complex with the Ubiquitin ligase E3 CUL3 protein, acting as a substrate-specific adaptor. Based on this, we named the gene as CUL3 substrate-adaptor protein 1 (*MtCSP1*).

To verify the interaction, a yeast two hybrid assay was performed by retransforming yeast with MtARFA1 and a clone that contains the complete ORF sequence of CSP1. To contrast the effect of the MtARFA1 activation state on the interaction, a mutant version of MtARFA1 in which the threonine 31 is exchanged by an asparagine, designated T31N, was also included in the assay. This mutation prevents the exchange of GDP for GTP, resulting in a DN conformation. No significant differences were observed between the different versions of MtARFA1, indicating that the interaction between MtCSP1 and MtARFA1 is independent of the MtARFA1 activation status (Figure 1A). Quantification of the interaction measuring β-Galactosidase activity showed that the strength of the interactions with MtCSP1 was similar between the CA and DN versions of ARFA1 (Figure 1B).

**Figure 1:**
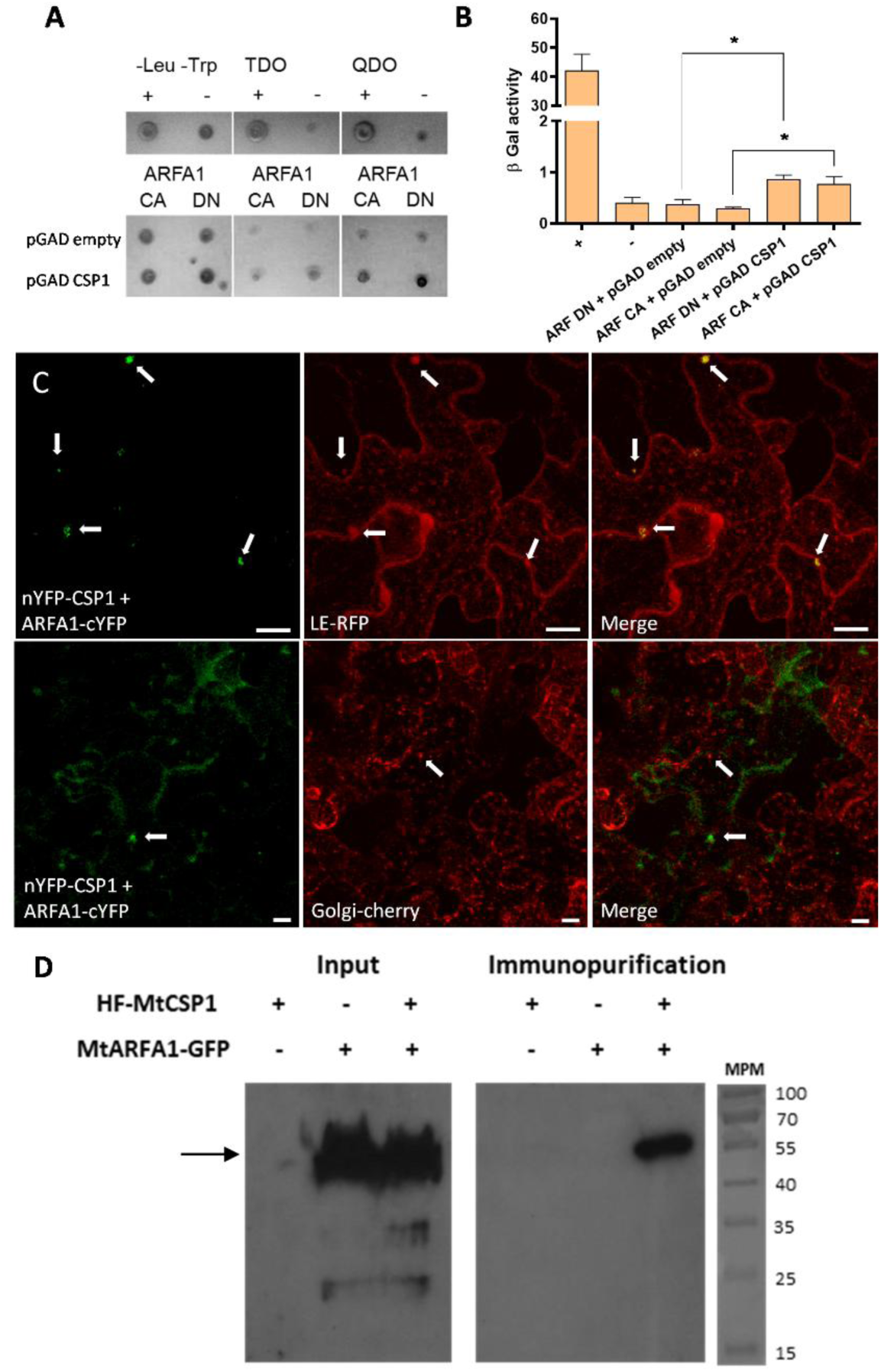
Interaction of MtARFA1 with MtCSP1. **A.** The ORFs of *MtARFA1* CA and DN forms were fused to the binding domain of the transcription factor Gal4 (BD) and introduced into the Y182 yeast strain, whereas the ORF of *MtCSP1* was fused to the activation domain of Gal4 (AD) and introduced into the AH109 yeast strain. Strains were mated and selected in triple or quadruple dropout medium (TDO and QDO, respectively). The empty vector pGADT7 was used as negative control (-), whereas the positive control was obtained by mating yeasts that carry the vectors pGADT7-Tag and pGBKT7-p53. **B.** β-galactosidase activity measured on the diploid yeasts using 3 replicates. **C.** Bimolecular fluorescence assay revealing the interaction of MtARFA1 and MtCSP1 (green) in *N. benthamiana* epidermal cells together with late endosome (LE) and Golgi markers (red). Arrows point to the fluorescent signal in small cytoplasmic vesicles. Bar = 25 μm. **D.** HIS-FLAG-MtCSP1 was transiently coexpressed with MtARFA1-GFP in *N. benthamiana* leaves. As negative controls, each construct was expressed alone. CoIPs were performed using Dynabeads coupled to antiFLAG antibodies (α-FLAG). Crude protein extracts (Input) and immunoprecipitated fractions were analyzed by immunoblotting using an anti-GFP antibody (α-GFP) to detect MtARFA1. The arrow indicates the expected size of MtARFA1-GFP.

To investigate the subcellular localization of the interaction between MtARFA1 and MtCSP1 and verify whether it occurs in plant cells, we performed a Bimolecular Fluorescence Complementation (BiFC) assay. The ORF of *MtCSP1* was fused to the N-terminal fragment of the split yellow fluorescent protein (nYFP-MtCSP1) and the ORF of *MtARFA1* was fused to the C-terminus fragment of the split YFP (MtARFA1-cYFP). These constructs were transformed in *Agrobacterium tumefaciens* and transiently expressed in *Nicotiana benthamiana* leaves. After two days, we observed YFP fluorescence in plant cells, particularly in small punctuated structures in the cytoplasm (Figure 1C, left panels). This result indicates that MtCSP1 interacts with MtARFA1 in vesicles, which is consistent with the subcellular localization previously reported for ARFA1 proteins (Pimpl *et al*., 2000; Takeuchi *et al*., 2002; Xu & Scheres, 2005; Stefano *et al*., 2006; Robinson *et al*., 2011). To further characterize the cellular punctuated structures where the MtARFA1-MtCSP1 interaction occurs, subcellular markers of late endosome (LE) and Golgi were used. We observed that the YFP signal corresponding to the MtARFA1-MtCSP1 interaction partially colocalized with the LE marker (Figure 1C, central top panel), whereas no colocalization was observed with the Golgi marker (Figure 1C, central low panel). The MtARFA1-MtCSP1 interaction was also tested in plant cells by co-immunopurification assays. A dual HIS-FLAG tag was fused to the N-terminal end of MtCSP1 (HF-MtCSP1), and the C-terminal end of MtARFA1 was fused to the green fluorescent protein (MtARFA1-GFP). These constructs were introduced in *N. benthamiana* leaves by agroinfiltration. Leaf tissue was used for immunopurification using magnetic beads coupled to α-FLAG antibody. The input and immunopurified fractions were analysed in an immunoblot assay revealed with α-GFP antibody (Figure 1D). A band of approximately 55 kDa corresponding to the estimated molecular weight of MtARFA1-GFP was observed only in the fraction immunopurified from leaves co-transformed with HF-MtCSP1 and MtARFA1-GFP, confirming the interaction between MtARFA1 and MtCSP1.

### Spatial expression pattern of *MtCSP1*

Publicly available expression data show that *MtCSP1* transcripts are present in several organs of *M. truncatula*, including seeds, flowers, leaves, roots and nodules. Interestingly, *MtCSP1* is strongly coregulated with *MtARFA1* along different organs and developmental stages (Supplemental Figure 2). To further characterize the expression of *MtCSP1*, we transformed *M. truncatula* roots with a plasmid containing the gene encoding the β-glucuronidase (GUS) under the control of a nearly 2 kb fragment of the *MtCSP1* promoter. GUS staining indicated that the promoter was active in nodules and in the tip of roots (Figures 2A and 2B, respectively). Activity was detected also during lateral root formation and emergence (Figure 2C). During symbiosis, *MtCSP1* promoter activity was detected in nodule primordia at the early stages of nodule development, as well as in the apical meristem and the lateral vasculature of developing nodules, whereas in mature nodules, the GUS activity was concentrated in the meristematic and differentiation zones (Figure 2D). These results indicate that the expression of *MtCSP1* is active in root meristems and during formation of the postembryonic root organs, lateral roots and nodules.

**Figure 2:**
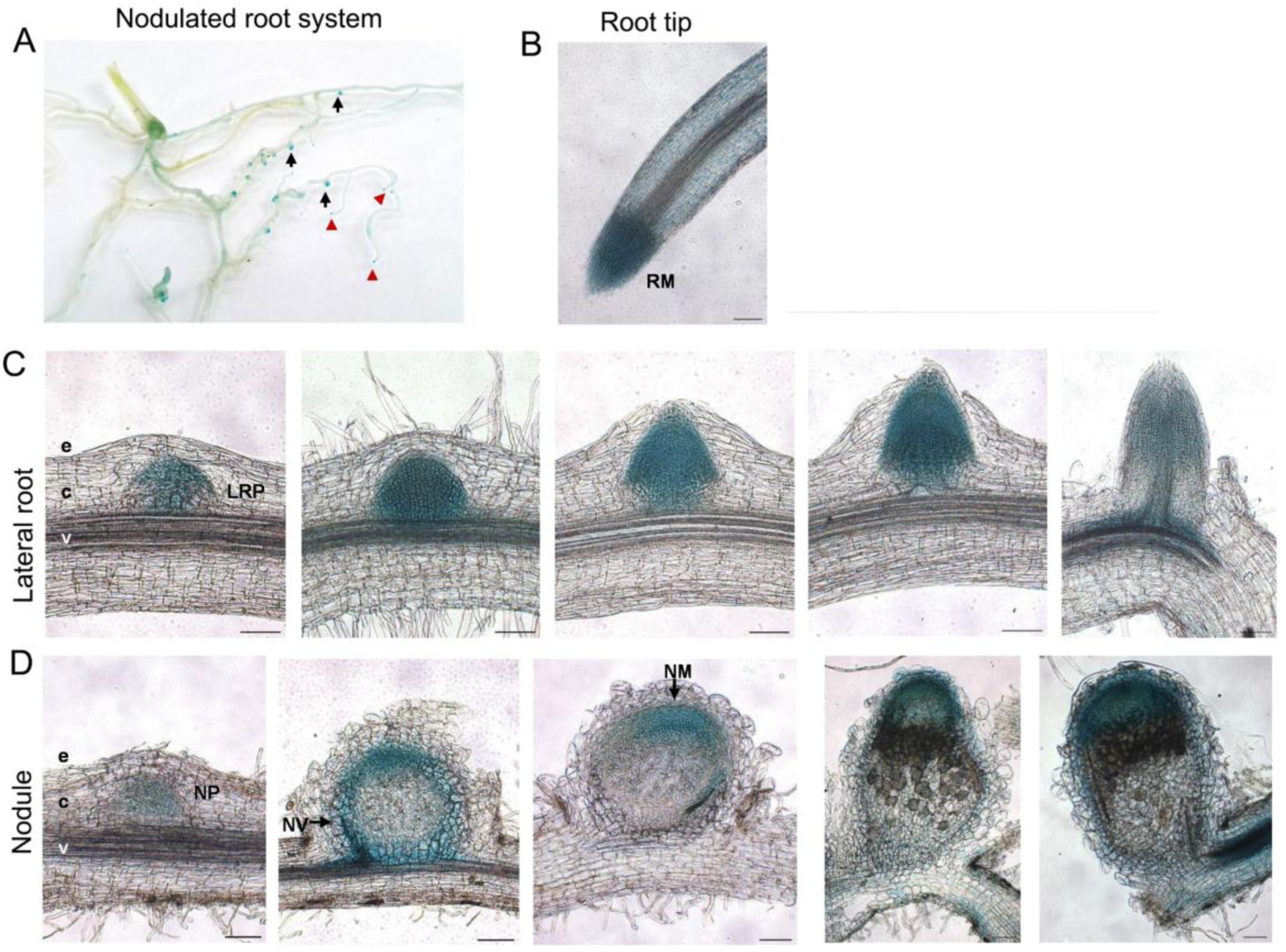
Activity of *MtCSP1* promoter in roots and nodules of *M. truncatula*. Histochemical GUS staining of roots and nodules transformed with the *pMtCSP1:GFP-GU*S construct. **A-B.** Transformed roots showing GUS activity in the meristematic zone of roots (red arrowheads) and nodules (black arrows). RM: Root Meristem. **C-D.** Spatiotemporal expression pattern of *MtCSP1* was analysed during lateral root (**C**) and nodule development (**D**). e: epidermis; c: cortex; v: vasculature; LRP: lateral root primordium; NP: nodule primordium; NV: nodule vasculature; NM: nodule meristem. Pictures are representative of images observed in three independent experiments each with more than 20 roots. Bars: 100 µm.

### Role of *MtCSP1* in nodulation

In order to functionally characterize *MtCSP1* in the nodulation context, we used *MtCSP1* RNA interference (RNAi)-mediated post-transcriptional silencing and overexpression strategies in roots obtained by *A. rhizogenes*-mediated transformation (Estrada-Navarrete *et al*., 2006). We amplified a 185 bp region of the 3’-UTR sequence of *MtCSP1* and cloned it as inverted repeats in the pK7GWIWG2D(II) vector (Karimi *et al*., 2002) under the control of the CaMV35S promoter to express an RNAi targeting *MtCSP1* (*CSP1* RNAi). Reverse transcription followed by quantitative PCR (RT-qPCR) analysis of transformed roots showed a marked and significant reduction in *MtCSP1* transcript levels as compared to *GUS* RNAi control roots (Supplemental Figure 3A). Composite plants transformed with either the *CSP1* RNAi or the *GUS* RNAi constructs were then inoculated with *S. meliloti*, and the number of nodules per root was quantified at different time points after inoculation. Nodulation kinetics showed a slight decrease in the number of nodules formed at early stages in the *MtCSP1* silenced roots as compared to the *GUS* RNAi control, with a significant reduction in mature nodules of 15- and 21-days post inoculation (dpi) (Figure 3A). This phenotype suggests that *MtCSP1* might be involved in the initiation of nodule formation. However, the size of the nodules was not significantly affected by expression of the *CSP1* RNAi (Figure 3B), indicating that nodule development was not significantly altered by silencing of *MtCSP1*. To analyze whether *MtCSP1* might have a role in the rhizobial infection, we quantified the number and progression of infection events by inoculating roots with a *S. meliloti* strain that expresses the Red Fluorescent Protein (RFP). The number of ITs per centimeter of root was not affected in *CSP1* RNAi roots at 7 dpi (Figure 3C). However, whereas many of the ITs formed in control roots were found to reach the dividing cells in the cortex, they ended more frequently in the root hair in *CSP1* RNAi roots (Figure 3D), indicating that the progression of ITs might be impaired or delayed by silencing of *MtCSP1*. As a complementary approach to characterize the function of *MtCSP1*, its ORF was fused to a double His-FLAG tag at the N-terminal and expressed under the control of the CaMV35S promoter (ox*MtCSP1*). *M. truncatula* composite plants were generated, and the overexpressing roots (Supplementary Figure 3B) were inoculated with *S. meliloti* and phenotypically characterized. It was observed that the number of nodules formed was higher when the *MtCSP1* gene was overexpressed as compared to control roots transformed with the empty vector (EV, Figure 4A). The size of nodules formed by roots overexpressing *MtCSP1* at 21dpi was similar to those of control plants (Figure 4B). On the other hand, there was no significant difference in the number of infection events (Figure 4D), but during the progression of infection events we observed a decrease in the infection events arrested at the microcolony stage, and an increase in the number of ITs that end in the root hair in oxCSP1 plants as compared to EV controls (Figure 4D). This shows that *MtCSP1* overexpression affects nodule organogenesis and progression of the ITs. Taken together, our results suggest that the *MtCSP1* is not required for initiation of ITs, but plays a key role in their progression and in nodule inception.

**Figure 3.**
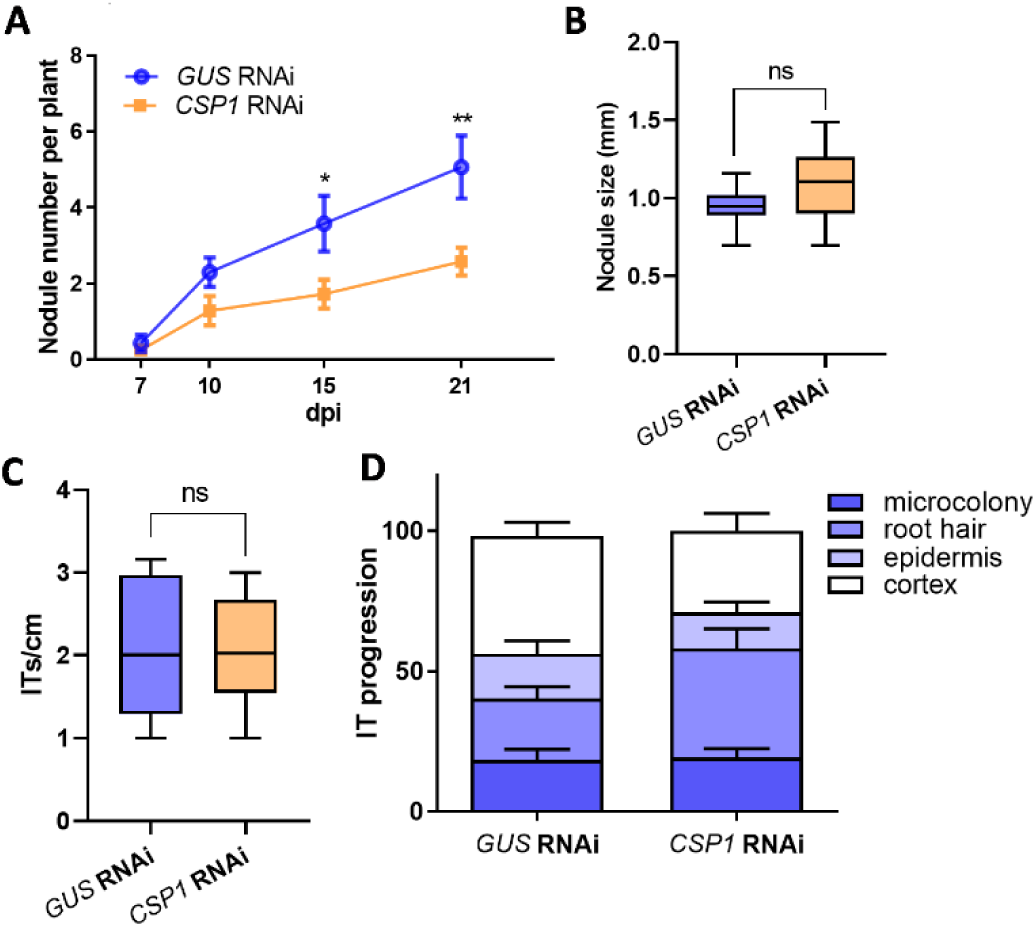
Symbiotic phenotypic analysis of *MtCSP1* silenced roots. **A.** Time-course nodule formation in *GUS* RNAi and *CSP1* RNAi roots upon inoculation with S*. meliloti*. Error bars represent mean ± SE of three independent biological replicates, each with at least 50 roots. **B.** Size of nodules formed in *GUS* and *CSP1* RNAi roots measured at 21dpi with *S. meliloti.* Each bar represents the mean ± SE of three biological replicates, each with more than 60 nodules. ns: not significant. **C.** Density of infection events in *GUS* and *CSP1* RNAi roots at 7dpi with a *S. meliloti* strain expressing the RFP protein. Each bar represents the mean ± SE of three biological replicates with more than 25 roots. ns: not significant. **D.** Progression of infection events in *GUS* and *CSP1* RNAi roots. Infection events were classified as microcolonies, ITs that end in the root hair, in the epidermal cell layer, or reach the cortex at 7 dpi. Each category is presented as the percentage of total infection events. Data are representative of three independent biological replicates with more than 25 transgenic roots. Asterisks indicate statistically significant differences in an unpaired two-tailed Student’s t-test (*, P ≤ 0.05; **, P ≤ 0.01).

**Figure 4.**
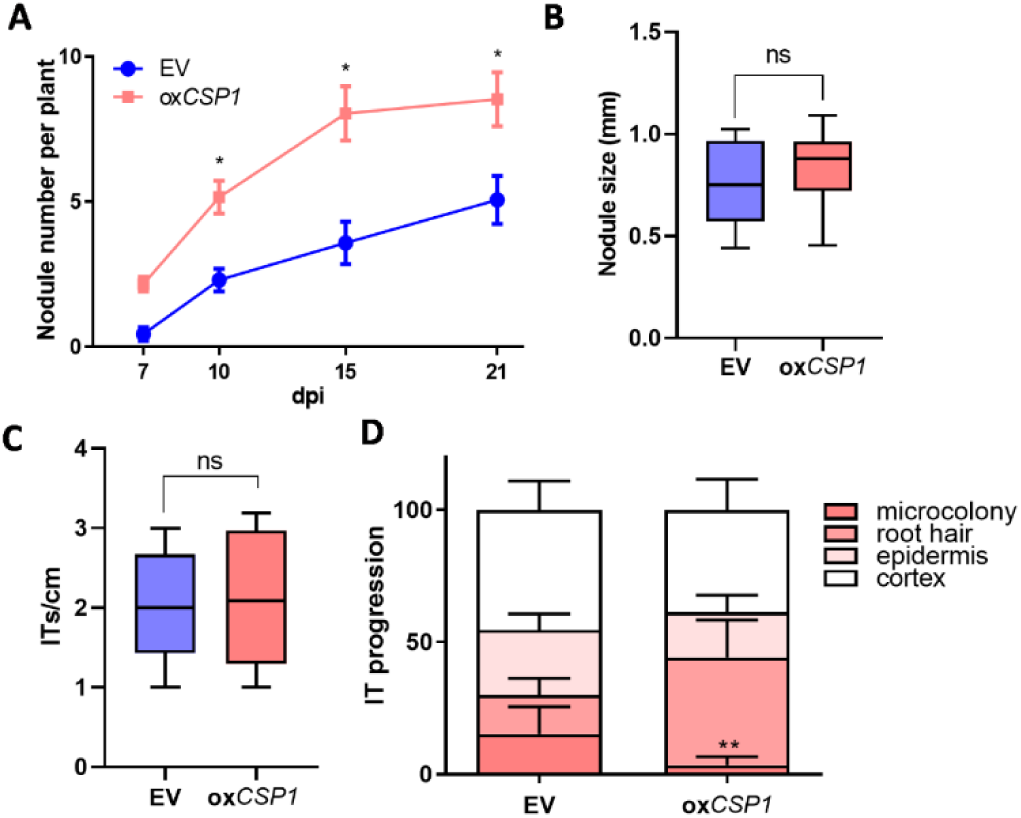
Symbiotic phenotypic analysis of *MtCSP1* overexpressing roots. **A.** Time-course nodule formation in EV and oxCSP1 roots upon inoculation with S*. meliloti*. Error bars represent mean ± SE of three independent biological replicates, each with at least 50 roots. **B.** Size of nodules formed in EV and oxCSP1 roots measured at 21dpi with *S. meliloti.* Each bar represents the mean ± SE of three biological replicates, each with more than 60 nodules. ns: not significant. **C.** Density of infection events in EV and oxCSP1 roots at 7dpi with a *S. meliloti* strain expressing the RFP protein. Each bar represents the mean ± SE of three biological replicates, each with more than 25 transgenic roots. ns: not significant. **D.** Progression of infection events in EV and oxCSP1 roots. Infection events were classified as microcolonies, ITs that end in the root hair, in the epidermal cell layer, or reach the cortex at 7 dpi. Each category is presented as the percentage of total infection events. Data are representative of three independent biological replicates, each with more than 25 transgenic roots. Asterisks indicate statistically significant differences in an unpaired two-tailed Student’s t-test (*, P ≤ 0.05).

### Expression of other components of the CUL3 complex

As mentioned in the introduction, BTB/POZ proteins are part of a complex involving CUL3 and RBX1 proteins. We wonder whether the expression of genes that encode these proteins is modulated during the symbiotic interaction with rhizobia. The *M. truncatula* genome has 8 genes with sequence similarity to the two Arabidopsis CUL 3 (Supplemental Figure 4A). Only two of them show detectable expression values in root postembryonic organs: one showed increased levels during lateral root formation and nodulation (MtrunA17Chr5g0416711), whereas the other did not show significant changes (MtrunA17Chr8g0361901) (Supplemental Figure 4B). Similarly, the four RBX1 genes found in the *M. truncatula* genome showed different expression patterns during the formation of lateral root organs. The gene MtrunA17Chr2g0322741, in particular, is strongly activated during lateral root formation and in response to rhizobium inoculation, whereas another member of the family, MtrunA17Chr7g0214441, is temporarily activated 10 hours after inoculation and then strongly repressed at later time points (Supplemental Figure 5). Altogether, these results show that mRNA levels of some members of the ubiquitin pathway involving the CUL3 ubiquitin ligase are modulated during formation of root post embryonic organs.

### Phylogenetic analysis of the BTB/POZ family in *M. truncatula*

To identify other genes with a BTB/POZ domain in *M. truncatula*, a genomic search was performed in the *Medicago truncatula* A17 r5.0 portal using the InterProscan program domain codes IPR000210 (BTB/POZ domain) and IPR003131 (Potassium channel tetramerization-type BTB domain), identified in the MtCSP1 sequence. A total of 58 genes were found that present at least one of these two codes (Supplemental Figure 6, Supplemental Table 1). To explore the phylogenetic relationships between the BTB/POZ-domain proteins, a multiple sequence alignment using the Clustal Omega tool, and a phylogenetic tree were generated using the FastTree software. These analyses revealed that the proteins in the clade in which MtCSP1 is found are highly divergent and exhibited little sequence conservation.

Besides BTB/POZ, CUL3 adaptor proteins contain additional domains that have been used to classify members of this protein family. As previously described in *Arabidopsis thaliana*, the most represented families are BTB NPH3, BTB only, BTB TAZ, BTB ankyrin and MATH BTB (Figure 5A) (Ban & Estelle, 2021). MtCSP1 is included in the group defined by the presence of a pentapeptide domain, a poorly explored subfamily that contains only one member in *A. thaliana* and *M. truncatula*. As previously described for other members of the complex, some of the genes encoding BTB proteins show differential accumulation during lateral root and nodule formation (Figure 5B). The transcripts encoding the two BTB/POZ proteins previously associated to determination of the nodule meristem, *NOOT1* and *NOOT2*, show an increase of their accumulation at early time points after rhizobia infection, whereas *MtCSP1* showed a milder response at 7 days after spot inoculation. Taken together, expression and genetic analyses suggest that, even though the tree genes are involved in nodule organogenesis, they play roles at different stages of the organ formation.

**Figure 5.**
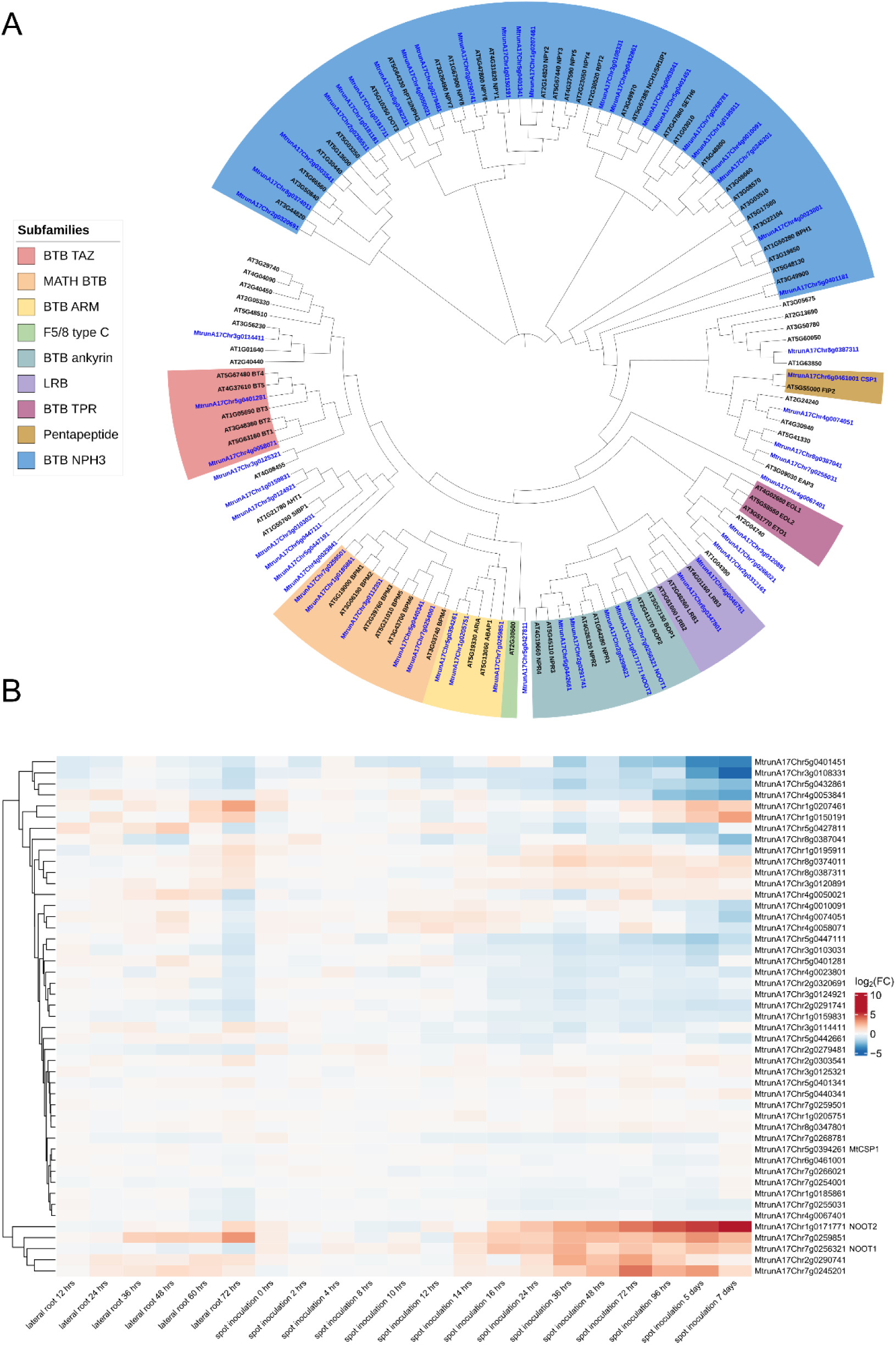
**A.** Phylogenetic analysis of BTB/POZ domain-containing proteins from *M. truncatula* and Arabidopsis. Protein sequence alignments generated with Clustal Omega were used to obtain the phylogenetic tree with FastTree v2.1.11. Numbers represent Bootstrap values from 1000 repeats. **B.** Expression of *MtCSP1* and other members of the BTB/POZ family using publicly available RNA-seq data from lateral root development and spot inoculation of rhizobia at different times. Data were obtained from Schiessl et al, 2019. Expression is represented as the log_2_ fold change relative to control plants without induction of lateral root formation (lateral root development) or mock-inoculated control plants (spot inoculation).

## DISCUSSION

The role of small GTPases in nodulation has been clearly established in previous studies; however, how the action of these molecular switches is regulated remains largely unknown. This work shows that a putative Cul3 substrate adaptor, MtCSP1, interacts with MtARFA1 in the context of the nitrogen-fixing symbiosis, establishing a connection between the genetic reprogramming of the root during the association with rhizobia and protein degradation via the ubiquitin pathway. *MtCSP1* has a positive effect on symbiosis as established by reverse genetics. The initial exchange of chemical signals between rhizobia and the root of legumes triggers two coordinated but independent genetic programs, one controlling the infection and the rhizobia progression through the IT, and the other triggering the formation of a new postembryonic organ, the nodule. Silencing by RNAi or overexpression of *MtCSP1* affected nodule formation and IT progression, supporting the hypothesis that this adaptor protein plays a role in the regulation of the both processes during nitrogen-fixing symbiosis. However, initiation of infection was not affected by *MtCSP1*, suggesting that the vesicle trafficking controlled by ARFA1 is necessary for growing of ITs, but not for their formation.

The interaction between monomeric GTPases and CUL3 substrate adaptors has been previously shown in animal systems. The leucine zipper–like transcriptional regulator 1 (LZTR1) binds to and targets members of the Ras family of GTPases for degradation, controlling cell proliferation (Bigenzahn *et al*., 2018; Steklov *et al*., 2018; Castel *et al*., 2019). Considering this, it is tempting to speculate that the role of *MtCSP1* could be associated with the degradation of MtARFA1 during the symbiotic interaction, once the GTPase has been activated to participate in the vesicle trafficking associated with symbiosis. A previous study showed that members of the Arabidopsis ARFA1 clade and other ARF proteins are targets of the ubiquitin protein degradation via the Arg/N degron pathway in Arabidopsis plants in the context of hypoxia responses (Zhang, H *et al*., 2024). Interestingly, a connection between hypoxia and nitrogen-fixing symbiosis has been proposed based on the oxygen depletion in the fixation zone required for nitrogenase activity (Pucciariello *et al*., 2019).

Several studies have established a connection between ubiquitination and nodulation (Hervé *et al*., 2011). It has been shown that the monomeric E3 ubiquitin ligase SIE3 interacts and ubiquitylates the Symbiosis Receptor Kinase (SymRK), a key regulator of the symbiotic process (Yuan *et al*., 2012). Like *MtCSP1*, overexpression of *SIE3* promoted nodulation, whereas downregulation by RNAi inhibited IT development and nodule organogenesis. Similarly, several E3 ligases were shown to interact and modulate levels of the Nod factors receptors, such as the Plant U-box protein 1 (MtPUB1) in *M. truncatula* (Mbengue *et al*., 2010), or the RING-type E3 ligase 1 (NIRE1) (Li *et al*., 2024) and Plant U-box proteins 7 and 13 in *L. japonicus* (Tsikou *et al*., 2018; Feng *et al*., 2025). These studies highlighted the crucial role of E3 ubiquitin ligases in controlling the root nodule symbiosis by modulating the levels of key components of the symbiotic signaling pathway by protein degradation prior and after rhizobium perception. Whereas these E3 ligases belong to the group of monomeric E3 proteins, CUL3 proteins form multimeric complexes with substrate adaptors such as MtCSP1. CUL3-adaptor proteins contain the BTB/POZ domain, composed by approximately 95 amino acids that form 5 alpha helixes and 3 beta strands, and other domains (e.g. armadillo, MATH, ankyrin, etc.) that define the specificity of substrate interactions and determine the subgroups of this family. The role of several substrate adaptors of this family has been previously established (Ferguson & Reid, 2005; Shen *et al*., 2019; Liu *et al*., 2023); however, pentapeptide-containing BTB/POZ is a branch of BTB/POZ with few members that has not been previously associated with root nodule symbiosis.

The interaction between MtARFA1 and MtCSP1 was detected in vesicles. We previously showed that the monomeric GTPAses RabA2 is localized in vesicles of the trans Golgi network and the LE, and they partially colocalize with ARFA1 in common bean (Blanco *et al*., 2009). These vesicles are concentrated at the tip of the root hair during IT formation, establishing a link between monomeric GTPases and early infection events (Peltzer Meschini *et al*., 2008). Interestingly, our results based on colocalization with subcellular markers showed that the interaction between MtCSP1 and MtARFA1 occurs in LEs, but not in the Golgi. MtCSP1, as part of the ubiquitination complex, is predicted to participate in the selection of the proteins that will be removed and delivered to the vacuole for degradation via the internalization of cell surface components and transport to the LE (Piper & Lehner, 2011). This removal of activated proteins can arrest or attenuate signaling processes mediated by receptors or molecular switches such as small GTPases, which determine the movement of cell material through vesicles. In this way, degradation of GTPases via the ubiquitination pathway can be important to control the progression of the genetic programs triggered by rhizobium.

One important question to understand the plasticity of the ubiquitin proteasome degradation system is to unveil how E3 ligases and their adaptors are expressed at specific developmental stages in certain tissues. Our results show that *MtCSP1* is expressed in the meristematic and infection region of mature nodules, as well as in young lateral roots. Publicly available expression data show a strong correlation between *MtARFA1* and *MtCSP1* accumulation, supporting the hypothesis that tight activation of the expression of *MtCSP1* might be required to mediate targeting of the MtARFA1 protein for degradation in mature nodules, once the infection is already established. Expression of other members of the CUL3-mediated ubiquitination system shows that some members of the CUL3, RBX1 and BTB/POZ substrate adaptors families changed their mRNA levels during development of lateral roots or nodules, suggesting that this particular type of cullin could be involved in these processes. Phylogenetic analysis indicates that the gene families of the CUL3 pathway are expanded in *M. truncatula* in comparison with Arabidopsis, a diversification that could be associated with biological processes occurring in legumes, such as the root-nodule symbiosis.

Control of gene expression includes protein turnover, allowing to degrade proteins that have been activated during specific stages of a biological process. The ubiquitin degradation pathway provides a versatile way to rapidly remove dispensable proteins contributing to the adaptation of plants to environmental changes or mediate developmental processes. Monomeric GTPases act as switches to turn on and off cellular processes such as vesicle trafficking and cytoskeleton rearrangements (Rivero *et al*., 2017). These processes are particularly active at early stages of symbiosis during changes of the root hair polar growth to form the IT and, later on, during release of bacteria and the formation of the symbiosomes. In addition to the interaction with ARFA1, MtCSP1 was previously shown to interact with formin (Banno & Chua, 2000), suggesting that it can also act as a CUL3 substrate adaptor of cytoskeletal proteins, which also play key roles during the development of ITs during infection. Future experiments will help to establish if MtARFA1 is ubiquitinated as a result of the interaction with MtCSP1 in the context of nodulation.

## EXPERIMENTAL PROCEDURES

### Biological material and plant transformation

*M. truncatula* WT Jemalong A17 seeds were obtained from INRA Montpellier, France (http://www.montpellier.inra.fr). DNA constructs were introduced in *Escherichia coli* DH5a*, A. rhizogenes* ARqua1 for hairy root transformations (Quandt, 1993), or *A. tumefaciens* GV3101 strains for *N. bethamiana* agroinfiltration assays. The *Sinorhizobium meliloti* strain 1021 (Meade, 1977) or the same strain expressing RFP (Tian *et al*., 2012) were used for root inoculation. All the strains were grown on LB (*E. coli*) or TY medium (*Agrobacterium*) supplemented with the appropriate antibiotic combinations. Plant growth and transformation were performed as previously described (Kirolinko *et al*., 2024).

#### Plasmid construction

To generate overexpression constructs, the open reading frame of *MtCSP1* was amplified by PCR using primers MtCSP1 OE F and MtCSP1 OE R (Supplementary Table S2), and cloned into the pENTR/D-TOPO vector (Invitrogen), creating pENTR-CSP1. For overexpression, the pENTR-CSP1 was recombined into the destination vector p35S:HF-GATA (Mustroph *et al*., 2009). For histochemical GUS staining assays, the 1978 bases upstream of the start codon of the *MtCSP1* genomic sequence were amplified with the specific primers promCSP1 F and promCSP1 R from Medicago genomic DNA, cloned in the pENTR/D-TOPO vector, and finally introduced by recombination into the destination vector pKGWFS7, driving the expression of the GFP-GUS fusion. For silencing of *MtCSP1* by RNAi, a 185 bp fragment corresponding to the 3′ UTR of *MtCSP1* was amplified by PCR, using CSP1 RNAi F and CSP1 RNAi R primers (Supplementary Table S2) and *M. truncatula* cDNA as a template. The PCR product was cloned into the entry vector pENTR/D-TOPO and recombined into the destination vector pK7GWIWG2D (II) (Karimi *et al*., 2002) to finally produce the *CSP1* RNAi construct.

#### Yeast two hybrid assays

The MtARFA1 Q71L (constitutively active, CA) and T31N (dominant negative, DN) mutated versions of MtARFA1 were obtained by directed mutagenesis as previously described for RABA2 (Dalla Via *et al*., 2017). The complete ORFs of mutated ARFA1 were obtained by PCR amplification using primers MtARFA1 *Eco*RI F and MtARFA1 *Sal*I R (Supplemental Table 2) and cloned into pGBKT7 using restriction enzymes *Eco*RI and *Sal*I. The resulting bait constructs were introduced into the Y187 (MATα) strain of *Saccharomyces cerevisiae*. To perform the double hybrid screening in yeast, competent cells of haploid yeast AH109 were transformed with a *M. truncatula* cDNA library fused to the Gal4 activation domain, obtained from total RNA of 4 dpi roots with *S. meliloti* strain 2011, and cloned into the pGAD-HA vector, kindly provided by the laboratory of Andreas Niebel (CNRS, Toulouse, France). Yeasts were mated with the haploid yeast strain Y187 containing the bait, resulting in diploid yeast carrying both constructs following instructions of the kit manufacturer (Clontech). The screening was performed on a total of 5×10^6^ diploid yeast cells. A suspension of 5×10^5^ cells/ml was inoculated at 200 µl per plate into 50 plates of 15 cm diameter. These were grown in QDO medium (without adenine, histidine, leucine and tryptophan) supplemented with 10mM 3-Amino-1,2,4-Triazole (3AT) (used as an inhibitor of the HIS3 reporter gene, known for its residual activity), for 7 days at 30°C. Plasmids were purified from yeast using the Yeastmaker Yeast Plasmid Isolation Kit (Clontech). The liquid β-galactosidase assay using *ortho*-Nitrophenyl-β-galactoside as substrate was performed according to the Yeast Protocols Handbook (Clontech).

#### Bimolecular fluorescence complementation

For BiFC assays, the ORFs of *MtCSP1* and *MtARFA1* were amplified by PCR using M13 primers from the corresponding pENTR/D-TOPO vectors and then recombined into the pGPTVII.Bar.YN-GW and pGPTVII.Hyg.GW-YC, respectively, for fusion to N termini of the N fragment of split YFP and the C terminus of the C fragment of split YFP, respectively (Hirsch *et al*., 2009). BiFC experiments were performed as previously described (Battaglia *et al*., 2014).

#### Co-inmunoprecipitation

A double HIS-FLAG tag was fused to the N-terminal end of MtCSP1 (HF-MtCSP1), and the C-terminal end of MtARFA1 was fused to the green fluorescent protein (MtARFA1-GFP) by LR recombination from the corresponding pENTR vectors into the p35S:HF (Mustroph *et al*., 2009) and pK7FWG2 (Karimi *et al*., 2002) Gateway vectors, respectively. Co-IP experiment was performed as previously described [44].

#### GUS staining

Histochemical GUS staining was performed as previously described (Rípodas *et al*., 2019). Roots were observed by bright field microscopy to visualize GUS staining in whole roots. Selected roots and nodules were embedded in 6% (w/v) agarose and cut to thin sections (70 μm) using a Leica VT1000 S Vibrating blade microtome. Tissue sections were analyzed by bright field microscopy in an inverted microscope OLYMPUS IX51.

#### Phenotypic analyses

For nodulation analysis, composite plants transformed with the *GUS* RNAi, *CSP1* RNAi, EV OE or ox*CSP1* constructs were transferred to slanted boxes containing nitrogen-free Fahraeus medium (Fahraeus, 1957) and inoculated with *S. meliloti* 1021 7 days after transplantation. The number of nodules was recorded at different time points after inoculation with *S. meliloti* as previously described (Hobecker *et al*., 2017). Nodule size was determined at 21 dpi with *S. meliloti* as previously described (Battaglia *et al*., 2014). For analysis of infection events, *GUS* and *CSP1* RNAi and EV OE and oxCSP1 plants of 14 days after transformation were transferred to petri dishes containing agar-Fahraeus medium free of nitrogen. Seven days after transplantation, roots were inoculated with the *S. meliloti* strain expressing RFP and grown as described (Kirolinko *et al*., 2021). Three independent biological replicates were performed for all experiments. In all cases, statistical significance of the differences for each parameter was determined by unpaired two-tailed Student’s t tests for each construct vs. the control lines.

#### RT-qPCR

RNA extraction, cDNA synthesis, and qPCR experiments were performed as described (Hobecker *et al*., 2017). Primers are listed in Supplementary Table 2.

#### Bioinformatics Analyses

Proteins with the BTB/POZ domain in *M. truncatula* were identified using InterProscan domain codes IPR000210 (BTB/POZ domain) and IPR003131 (Potassium channel tetramerization-type BTB domain) in the INRA (National Institute for Agricultural Research) genomic portal “Medicago truncatula A17 r5.0 genome portal” (https://medicago.toulouse.inra.fr/).

Candidate genes for Cullins and RBX1 in the *M. truncatula* genome were identified using BLASTP searches with the sequences of proteins previously described and classified in *A. thaliana* (Gray *et al*., 2002; Risseeuw *et al*., 2003). The list of identified genes was curated and classified manually.

Protein sequences were aligned using Clustal Omega (Sievers *et al*., 2011) via the online platform NGPhylogeny.fr (Lemoine *et al*., 2019). Multiple alignments were performed using a full distance matrix, and five iterations were performed to refine the alignment quality. Phylogenetic trees were constructed using FastTree v2.1.11 (Price *et al*., 2009; Price *et al*., 2010) with the LG substitution model (Le & Gascuel, 2008). An approximate maximum likelihood method was applied, and branch robustness was assessed using 1000 Bootstrap replicates. The resulting trees were visualized and annotated using iTOL (Interactive Tree of Life) (Letunic & Bork, 2024).

Gene expression was explored using publicly available datasets. Lateral root and node expression data at different time points were obtained from (Schiessl *et al*., 2019). Heatmaps were generated in R-studio (RStudio Team (2019). RStudio:

Integrated Development for R. RStudio, Inc., Boston, MA URL, http://www.rstudio.com/) using the pheatmap library. In all cases, hierarchical clustering of rows was performed using Euclidean distance and average linkage method.

## ACKNOWLEDGEMENTS

We would like to thank Andreas Niebel for providing the yeast two-hybrid library. CRip, FAB and MEZ are members of CONICET. MC, AE and CRiv were funded by CONICET fellowships.

## FUNDING

This work was financially supported by grants from ANPCyT, Argentina (PICT 2019/00029, 2020-00053 and 2021-00170), Secretaría de Ciencia, Tecnologıa e Innovacion of Argentina (RIBOLEG, CONVE-2023-100766842) and by the International Research Project LOCOSYM of the CNRS.

## AUTHOR CONTRIBUTION

All authors designed experiments and analyzed the data; CRip and CRiv performed interaction and phenotypic assays, MC developed expression analysis and AE was in charge of *in silico* analyses. MEZ and FAB conceived research and supervised the study.

## DECLARATION OF INTERESTS

The authors declare no competing interests.

## SUPPLEMENTAL INFORMATION

**Supplemental Figure 1.** Coexpression of *MtRABA2* and three members of the *M. truncatula* ARFA1 subfamily.

**Supplemental Figure 2.** Transcript levels of *MtCSP1* and *MtARFA1* in public datasets.

**Supplemental Figure 3.** Expression levels of *MtCSP1* in *CSP1* RNAi and ox*CSP1* roots.

**Supplemental Figure 4.** Phylogenetic tree and gene expression of cullin proteins from Arabidopsis and *M. truncatula*.

**Supplemental figure 5.** Phylogenetic tree and gene expression of RBX1 proteins from Arabidopsis and *M. truncatula*.

**Supplemental figure 6.** Localization of domains in the BTB/POZ proteins of *M. truncatula*.

**Supplemental Table 1.** Members of the BTB/POZ family of substrate adaptors in *M. truncatula*.

**Supplemental Table 2.** List and sequences of primers used in this study.

## Notes

### Competing Interest Statement

The authors have declared no competing interest.

