## Supplementary information for "The GTPase ARFA1 interactor Cullin 3 Substrate-adaptor Protein 1 (CSP1) positively modulates nodulation"

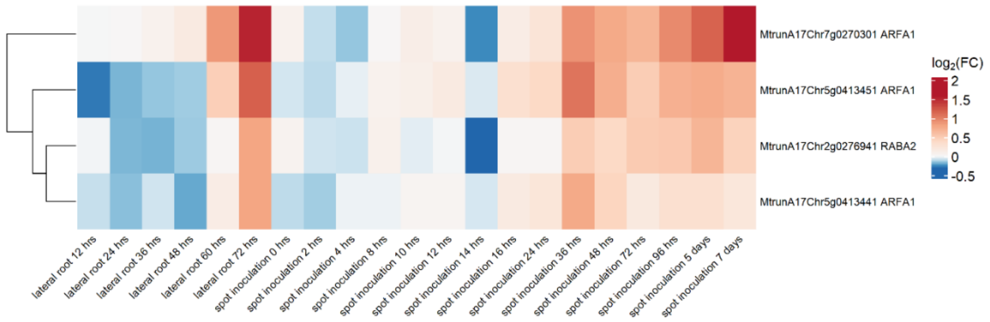


**Supplemental Figure 1.** Coexpression of *MtRABA2* and three members of the *M. truncatula* ARFA1 subfamily using data from lateral root development and spot inoculation of rhizobia at different times. Data were obtained from Schiessl et al, 2019. Expression is represented as the $log_2$ fold change relative to control plants without induction of lateral root formation (lateral root development) or mock-inoculated control plants (spot inoculation).

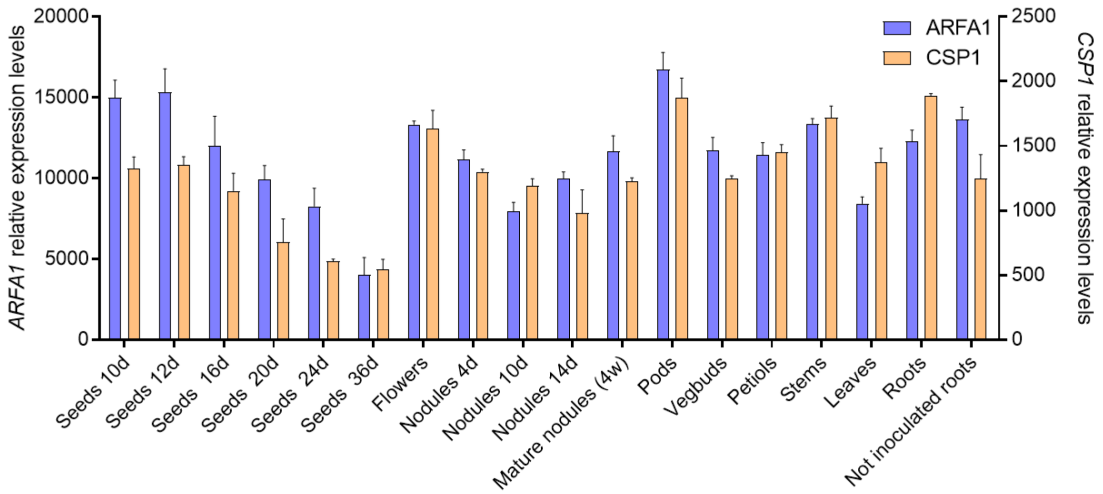


**Supplemental Figure 2.** Transcript levels of *MtCSP1* and *MtARFA1* in seeds of 10, 12, 16, 20, 24 and 36 days after pollination, flowers, nodules at 4, 10 and 14 days post inoculation (Nod 4dpi, Nod 10dpi, Nod 14dpi), mature nodules of 4 weeks post inoculation, pods, vegetative buds (Vegbuds), petioles, stems, leaves, 28-day old roots (Roots) and uninoculated roots. Microarray data reported by Benedito et al (2008) were obtained from the *Medicago truncatula* Gene Expression Atlas (MtGEA), publicly available in https://mtgea.noble.org/v3/ (Carrere et al, 2021).

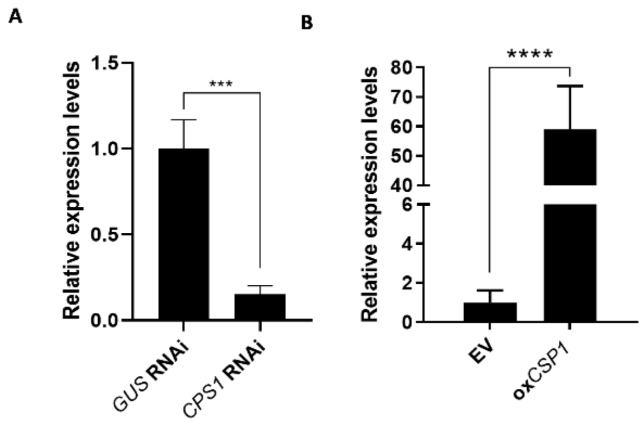


**Supplemental Figure 3.** Transcript levels of *MtCSP1* in *CSP1* RNAi (**A**) and ox*CSP1* roots (**B**). Expression values were determined by RT-qPCR with primers specific for *MtCSP1*. Values were normalized to *MtHIS3L* and expressed relative to the *GUS* RNAi or to the empty vector (EV) samples, which were set at 1. Each bar represents the mean ± SE of three biological replicates. Asterisks indicate statistically significant differences between *GUS* and *CSP1* RNAi roots in an unpaired two-tailed Student's t-test (*** $P \le 0.005$; **** $P \le 0.001$).

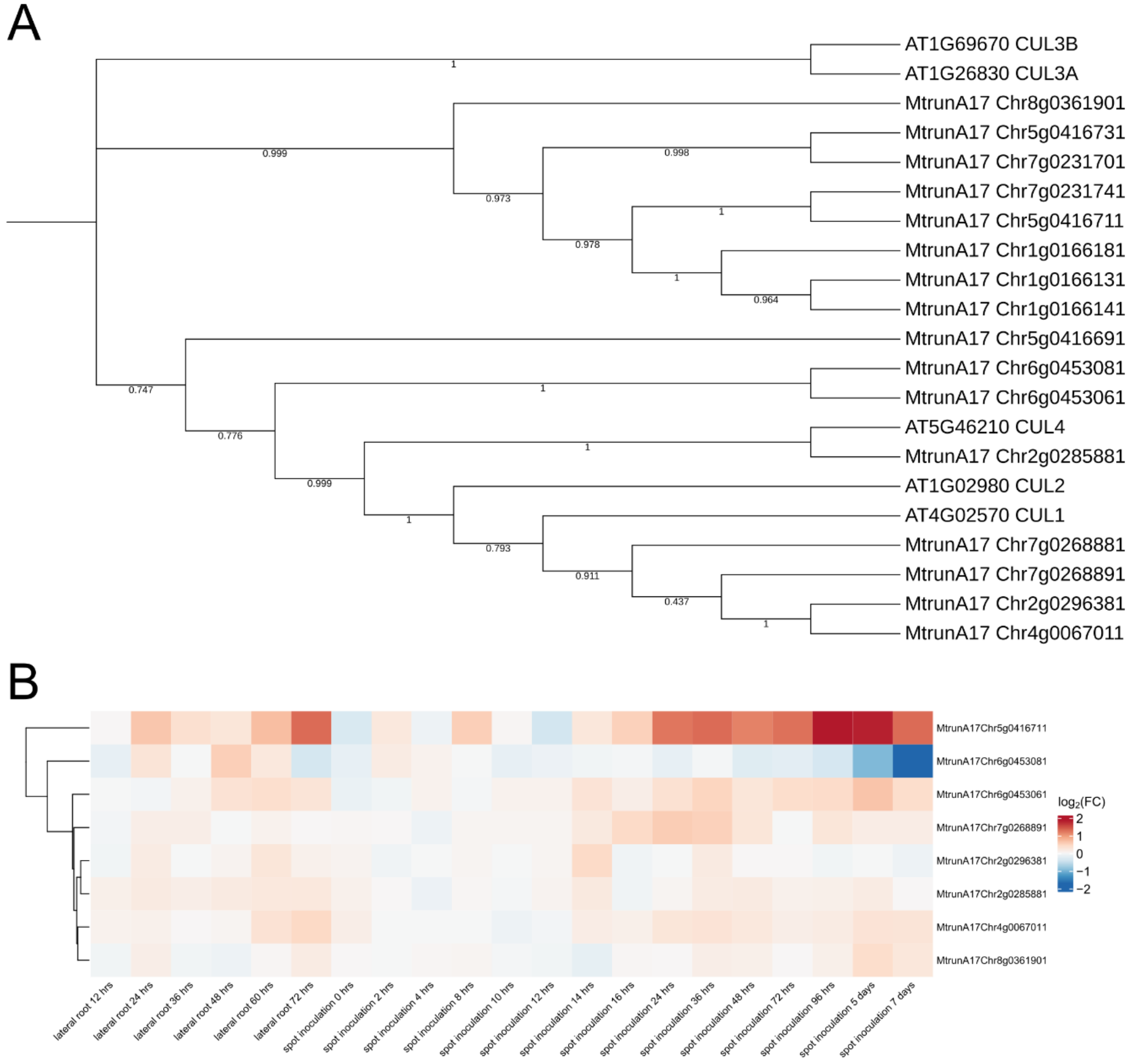


**Supplemental Figure 4. A.** Phylogenetic tree showing cullin proteins from Arabidopsis and *M. truncatula*. **B.** Expression of *M. truncatula* cullin genes using data from lateral root development and spot inoculation of rhizobia at different time points. Data were obtained from Schiessl et al, 2019. The heatmap shows genes with detectable transcript levels under all conditions. Expression is represented as the $\log_2$ fold change relative to control plants without induction of lateral root formation (lateral root development) or mock-inoculated control plants (spot inoculation).

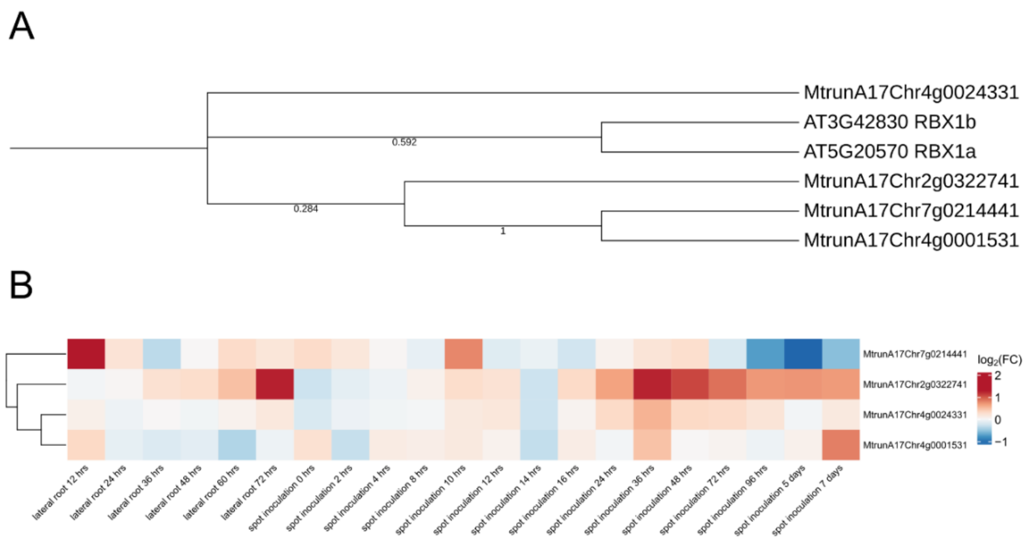


**Supplemental figure 5.** Phylogenetic tree showing RBX1 proteins from Arabidopsis and *M. truncatula*. **B.** Expression of *M. truncatula* homologs of RBX1 using data from lateral root development and spot inoculation of rhizobia at different times. Data were obtained from Schiessl et al, 2019. Expression is represented as the $log_2$ fold change relative to control plants without induction of lateral root formation (lateral root development) or mock-inoculated control plants (spot inoculation).

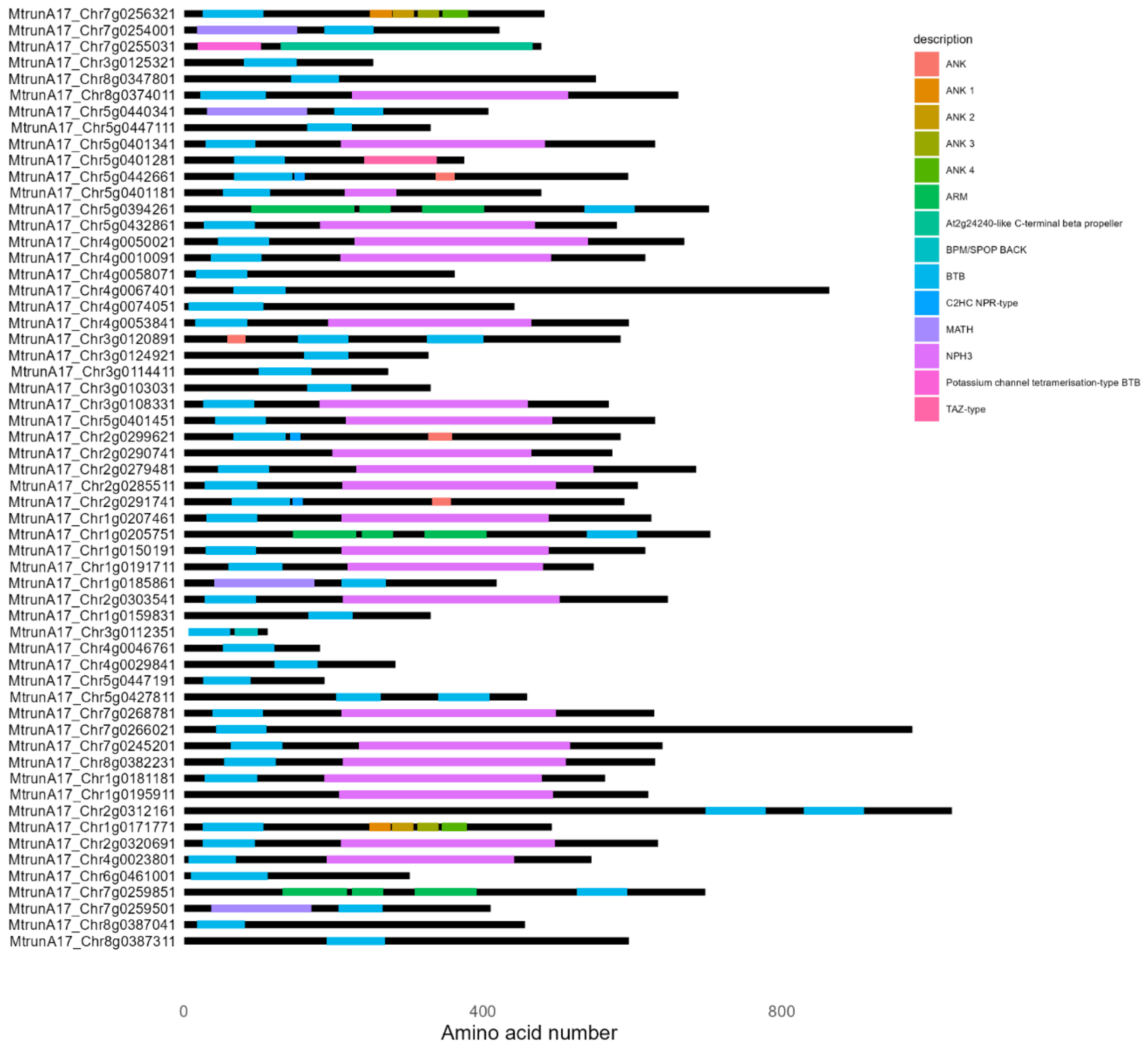


**Supplemental Figure 6.** Structure of *M. truncatula* BTB/POZ proteins. X axis represents the length of amino acid sequences. Colored boxes represent conserved domains and regions. Data was obtained from Uniprot protein database and plotted using the drawProteins package in R.

**Supplemental Table 2.** List and sequences of primers used in this study.

| Primer | Sequence |
|---|---|
| MtCSP1 OE F | CACCATGAAGGAATTCCAATCCGA |
| MtCSP1 OE R | TTATCTGATTGCTCCGAGCA |
| CSP1 RNAi F | CACCCAAGTGGGGATGTGGTTAAAG |
| CSP1 RNAi R | GTAAGGAGGAAGCCATTTAGG |
| qCSP1 F | CAAGTGGGGATGTGGTTAAAG |
| qCSP1 R | CACCTAAATGGCTTCCTCCTTAC |
| PromCSP1 F | CACCGAAGTTGTGTTGGTAGTTGG |
| PromCSP1 R | TGTAGGAGAAATAACCTTACTCA |
| MtARFA1 EcoRI F | CGGAATTCGGATTGTCATTCACGAAGCT |
| MtARFA1 SalI R | CGACGTCGACCTATGCCTTGTTTGCAATGT |
